# Simultaneous profiling of multiple chromatin proteins in the same cells

**DOI:** 10.1101/2021.04.27.441642

**Authors:** Sneha Gopalan, Yuqing Wang, Nicholas W. Harper, Manuel Garber, Thomas G. Fazzio

## Abstract

Methods derived from CUT&RUN and CUT&Tag enable genome-wide mapping of the localization of proteins on chromatin from as few as one cell. These and other mapping approaches focus on one protein at a time, preventing direct measurements of colocalization of different chromatin proteins in the same cells and requiring prioritization of targets where samples are limiting. Here we describe multi-CUT&Tag, an adaptation of CUT&Tag that overcomes these hurdles by using antibody-specific barcodes to simultaneously map multiple proteins in the same cells. Highly specific multi-CUT&Tag maps of histone marks and RNA Polymerase II uncovered sites of co-localization in the same cells, active and repressed genes, and candidate cis-regulatory elements. Single-cell multi-CUT&Tag profiling facilitated identification of distinct cell types from a mixed population and characterization of cell type-specific chromatin architecture. In sum, multi-CUT&Tag increases the information content per cell of epigenomic maps, facilitating direct analysis of the interplay of different proteins on chromatin.

## Introduction

Regulation of gene expression is a complex process involving integration of numerous positive and negative inputs from multiple regulatory proteins on a small number of cis regulatory elements (CREs) for each gene (Bulger and Groudine, 2011; Ong and Corces, 2011). CREs control the recruitment and activation of RNA Polymerase II (RNAPII) to control the production of RNA transcripts. Many different epigenetic regulators combine to determine CRE activity, including histone modifying enzymes, nucleosome remodeling factors, and enzymes that add covalent modifications to DNA bases. In turn, these epigenetic marks largely control binding of transcription factors and recruitment of additional chromatin remodeling proteins to modulate recruitment of RNAPII.

Although the identities of most regulatory proteins and epigenetic marks are now known, the mechanisms by which their regulatory inputs are integrated to produce a broad range of transcript levels remain poorly understood. For nearly 15 years, chromatin immunoprecipitation followed by deep sequencing (ChIP-seq) has been widely used for genome-wide mapping of proteins on chromatin (Gilmour and Lis, 1984; Park, 2009). However, numerous technical limitations limit the utility of ChIP-seq, particularly under conditions where samples are limiting. Two newer sets of methods named CUT&RUN (Skene and Henikoff, 2017) (along with similar methods (Hainer et al., 2019; Janssens et al., 2018; Ku et al., 2019)), and CUT&Tag (Kaya-Okur et al., 2019) (and related methods (Carter et al., 2019; Wang et al., 2019)) have greatly improved the sensitivity and specificity of chromatin profiling. Each of these methods utilizes enzymatic release and subsequent sequencing of small chromatin fragments surrounding the protein of interest. As a result, nonspecific chromatin is not fragmented and subjected to immunoprecipitation, vastly reducing background and producing libraries with high signal-to-noise that require low sequencing depth.

Despite these substantial improvements in chromatin mapping, CUT&RUN and CUT&Tag do not provide insights into how various regulatory proteins function together on CREs to dictate transcriptional output. As with ChIP-seq, these mapping approaches profile one protein at a time, requiring comparisons of multiple different maps to infer the overall composition of chromatin domains. However, since different chromatin proteins or histone marks are mapped in different samples of cells, one cannot distinguish co-binding of two different proteins in the same cells from alternative binding of one protein or the other (but not both) at the same sites. This is a particular concern for CREs, where dozens of regulatory proteins (some of which are present within protein complexes measuring over one megadalton in mass) all map to the same locations, despite the potential steric problems associated with co-binding of all these proteins. An alternate approach is to perform sequential ChIP-seq (with multiple variations, including Re-ChIP and co-ChIP), where two different antibodies are used sequentially to immunoprecipitate DNA co-bound by two chromatin proteins or histone marks (Geisberg and Struhl, 2004; Kinkley et al., 2016; Weiner et al., 2016). However, sequential ChIP-seq protocols typically capture only the overlapping locations of chromatin proteins, rather than the complete genomic maps of each. In addition, these methods can be difficult to implement for many chromatin proteins, as the need for multiple immunoprecipitation steps compounds many of the inefficiencies of ChIP-seq. Consequently, it can be difficult to adapt this approach for small or heterogenous populations of cells.

Here, we developed a method based on the CUT&Tag (Kaya-Okur et al., 2019) approach that enables simultaneous mapping of multiple chromatin proteins or histone modifications in the same cells, which we denote “multi-CUT&Tag”. We use protein A-Tn5 transposase (pA-Tn5) loaded with antibody-specific barcodes to simultaneously treat cells with multiple different antibodies, each recognizing a different protein or epitope on chromatin. We find that multi-CUT&Tag exhibits high sensitivity and specificity, similar to that of standard CUT&Tag, and produces maps of different chromatin proteins that are concordant with orthogonal ChIP-seq maps of the same proteins. Multi-CUT&Tag maps revealed coassociation of RNAPII with histone modifications, as well as co-association of active and repressive histone modifications at numerous sites, validating this approach as a probe of coassociation of chromatin proteins. We further show that multi-CUT&Tag can be adapted for single-cell profiling in high throughput. Single-cell multi-CUT&Tag profiling of repressive and activating histone marks H3K27me3 and H3K27ac enabled clustering of cell types within a mixed population, and inference of cell type-specific chromatin architecture. Together, these studies demonstrate that multi-CUT&Tag overcomes several limitations of current chromatin profiling methodologies by simultaneously identifying the genome-wide binding sites of multiple proteins in the same cells.

### Design

Multi-CUT&Tag was designed to fill two voids in the current toolkit of chromatin mapping technologies—the inability to profile multiple proteins from the same samples, necessitating prioritization of one or a few epitopes when samples are limiting, and the inability to directly measure co-association of chromatin proteins in cells. To this end, we developed a strategy to modify the CUT&Tag approach such that pA-Tn5 carries and deploys adapters with barcodes that are unique to different antibodies. Such an approach would allow different antibodies to be used in the same cells, enabling simultaneous mapping of multiple chromatin-associated proteins or histone modifications. This strategy, which we denote “multi-CUT&Tag”, has several unique features (Figure 1A), described below.

**Figure 1:**
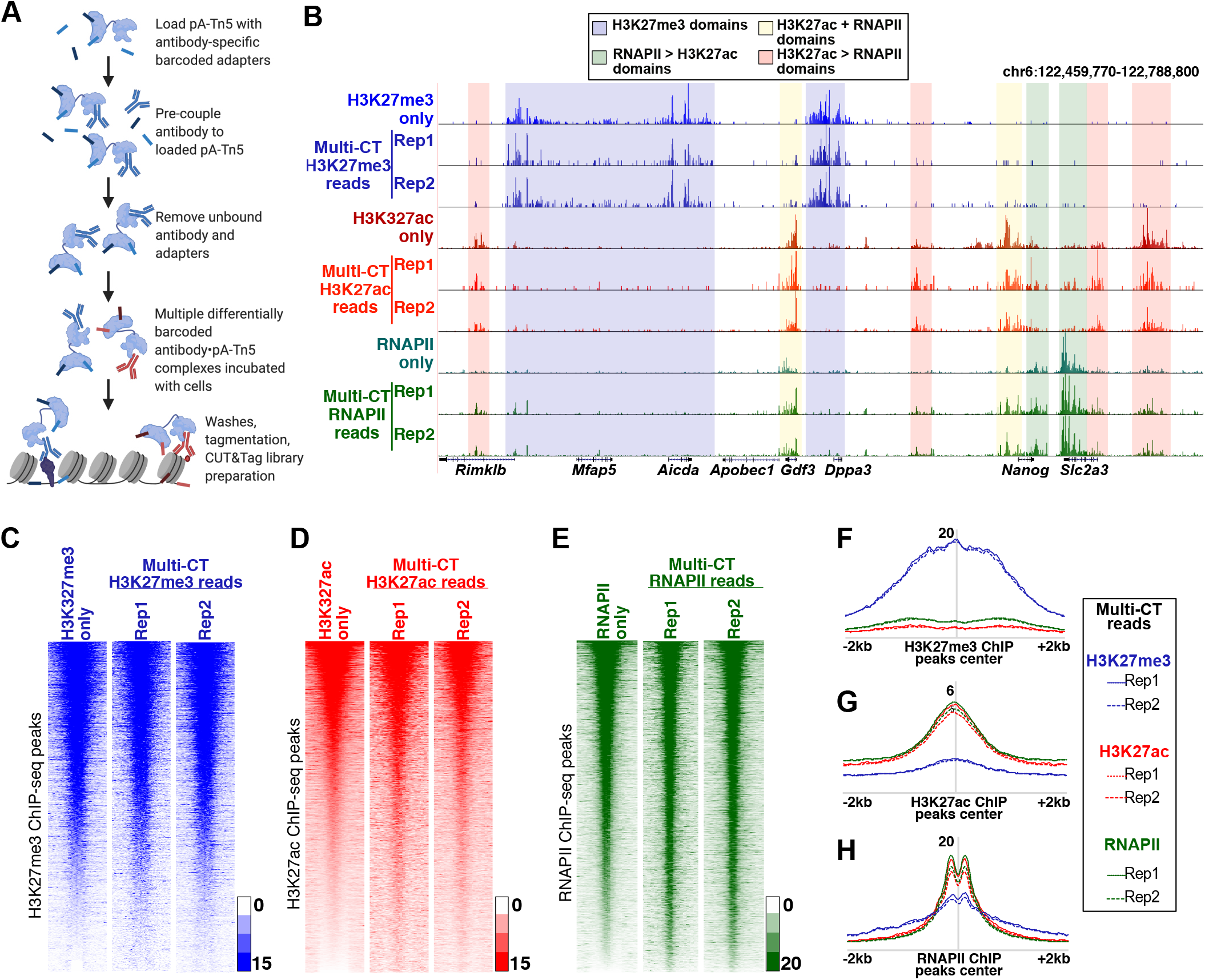
Simultaneous profiling of multiple chromatin proteins using multi-CUT&Tag. **A,** Diagrammatic representation of multi-CUT&Tag workflow. **B,** Genomic landscape comparing single Ab CUT&Tag and triple Ab multi-CUT&Tag for H3K27me3, H3K27ac and RNAPII. The shaded regions represent domains with high enrichment of the indicated proteins or histone modifications. **C-E,** Heatmaps depicting enrichment of reads from single Ab CUT&Tag and triple Ab multi-CUT&Tag over the genomic locations of peaks of published ChIP-seq data specific for H3K27me3 (+/-5kb; GSE123174) (C), H3K27ac (+/-2kb; GSE31039) (D) and RNAPII (+/-2kb; GSE112113) (E). **F-H**, Average enrichment of reads from triple Ab multi-CUT&Tag over the peak locations (called from published ChIP-seq data) of H3K27me3 (F), H3K27ac (G) and RNAPII (H).

First, we constructed and purified a version of pA-Tn5 with an N-terminal 6-histidine (6-His) tag to facilitate purification after pA-Tn5 is loaded with oligonucleotide adapters and precoupled to antibodies (Abs). We used several barcoded Tn5 adapters described previously (Amini et al., 2014). After adapter loading, an approximately two-fold excess of barcoded pA-Tn5 protein is incubated with an antibody of interest, to form an antibody•pA-Tn5 complex (Ab•pA-Tn5). Next, uncomplexed antibody and free adapters are removed by binding pA-Tn5 to TALON beads (which binds the 6-His tag on pA-Tn5), followed by elution of Ab•pA-Tn5 with imidazole, and subsequent buffer exchange. This strategy enables reads with barcodes specific to each Ab to be directly assigned to that Ab and segregated from reads with different barcodes to generate Ab-specific maps, as described below. Conjugates corresponding to different antibodies are then used in desired combinations for mapping, using a series of incubation and washing steps similar to the original CUT&Tag approach (Kaya-Okur et al., 2019). After sample cleanup, fragments are PCR amplified to attach a second (sample-specific) barcode and the P5 and P7 elements necessary for sequencing on the Illumina platform.

Due to the presence of Ab-specific barcodes, we required a custom sequencing and demultiplexing strategy for multi-CUT&Tag libraries (Figure S1A). Reads were sequenced using custom sequencing oligos that first read through the Ab-specific barcodes, followed by the mosaic-end sequence common to all Tn5 adapters, and finally read into the genomic loci targeted by pA-Tn5. Indexing cycles using custom indexing primers were used to identify different samples. Libraries were demultiplexed based on sample-barcodes and further segregated based on Ab-specific barcodes, followed by trimming of Ab-barcodes and adapterspecific sequences. The resulting (trimmed) reads were then aligned to the genome. This pipeline, along with a few key steps of the library generation strategy, were slightly modified for single cell sequencing, as described in the Methods.

In addition to generating specific maps of different chromatin proteins from the same cells, multi-CUT&Tag was designed to directly detect colocalization of different proteins at the same genomic location. In cases where two proteins co-localize, two distinct Ab•pA-Tn5 complexes will be present within close proximity on the chromosome and can each insert a differentially barcoded set of adapters into nearby DNA. Some reads in this scenario will harbor “mixed” barcodes—different antibody barcodes at each end. These reads are identified in the process of demultiplexing, and can be compared to maps derived from homogeneous reads from each single Ab. In sum, multi-CUT&Tag is designed to leverage the benefits of “standard” CUT&Tag approaches (e.g., high sensitivity and ease-of-use) in a unique strategy that enables direct detection of proteins co-bound to the same sites in the same cells, as well as reducing the sample requirements and hands-on workload of epigenomic profiling studies.

## Results

### Simultaneous mapping of multiple chromatin proteins

To test this approach, we used various combinations of Ab•pA-Tn5 complexes to map the genomic locations of two histone modifications and the elongating form of RNA Polymerase II in mouse embryonic stem cells (mESCs). We examined H3K27me3 (a mark of repressed genes and CREs), H3K27ac (a mark of active genes and CREs), and RNA polymerase II phosphorylated on serine 2 of its C-terminal domain (a mark of actively elongating RNA polymerase II; hereafter denoted simply as RNAPII).

We tested the three individual Ab•pA-Tn5 conjugates alone, in all pair-wise combinations, and with the three Ab•pA-Tn5 conjugates combined, in pools of 100,000 mESCs. The reads from each multi-CUT&Tag library were segregated based on the antibody barcode, and individual maps specific for H3K27me3, H3K27ac, or RNAPII were generated as described in the Methods. Significantly, for all three two Ab multi-CUT&Tag combinations (Figure S1B-D) and the three Ab multi-CUT&Tag combination (Figure 1B), we found that the genomic landscapes of each epitope were nearly identical to those observed in single antibody control experiments. As an orthogonal measure, we examined enrichment of multi-CUT&Tag maps for each epitope at peaks called from published ChIP-seq data from other sources (Mu et al., 2018; Zhang et al., 2018), finding that both single Ab and multi-CUT&Tag libraries were highly concordant with ChIP-seq maps of the same epitopes (Figure 1C-H; Figure S2A-C).

### High sensitivity and specificity of multi-CUT&Tag profiles

To gain a quantitative measure of the specificity and reproducibility of multi-CUT&Tag experiments, we measured the correlation among multi-CUT&Tag maps from different Abs, as described in the Methods. For each individual epitope, we observed very high correlation between single Ab maps and maps from multi-CUT&Tag experiments, demonstrating high reproducibility (Figure 2A). In addition, we observed moderate to strong correlation between H3K27ac and RNAPII maps from both single Ab and multi-CUT&Tag libraries, consistent with their established roles in gene expression. Conversely H3K27ac/RNAPII maps correlated poorly with maps of H3K27me3, a repressive mark, as expected (Figure 2A). Reads corresponding to all three epitopes were enriched near gene TSSs (Figure S2D-E), as anticipated from their established patterns of localization. However, TSSs most strongly enriched for H3K27ac/RNAPII were generally poorly enriched for H3K27me3, and vice versa (Figure S2F). These results demonstrate the specificity of multi-CUT&Tag maps. Furthermore, these findings confirm that multi-CUT&Tag can not only generate accurate maps of proteins that are largely non-overlapping (e.g., H3K27me3 and RNAPII), but can also simultaneously map epitopes that strongly overlap throughout the genome (e.g., H3K27ac and RNAPII). As an additional measure of specificity, we examined the fraction of reads in peaks (FRIP), a simple metric for signal to noise that has been adopted widely in genomic mapping studies (Landt et al., 2012). We observed FRIP scores from 0.35-0.70 for multi-CUT&Tag libraries for the three antibodies tested (Figure S2G-I), consistent with the high specificity observed in our single Ab maps and traditional CUT&Tag protocols (Kaya-Okur et al., 2019). These data confirm the high specificity and reproducibility of multi-CUT&Tag.

**Figure 2:**
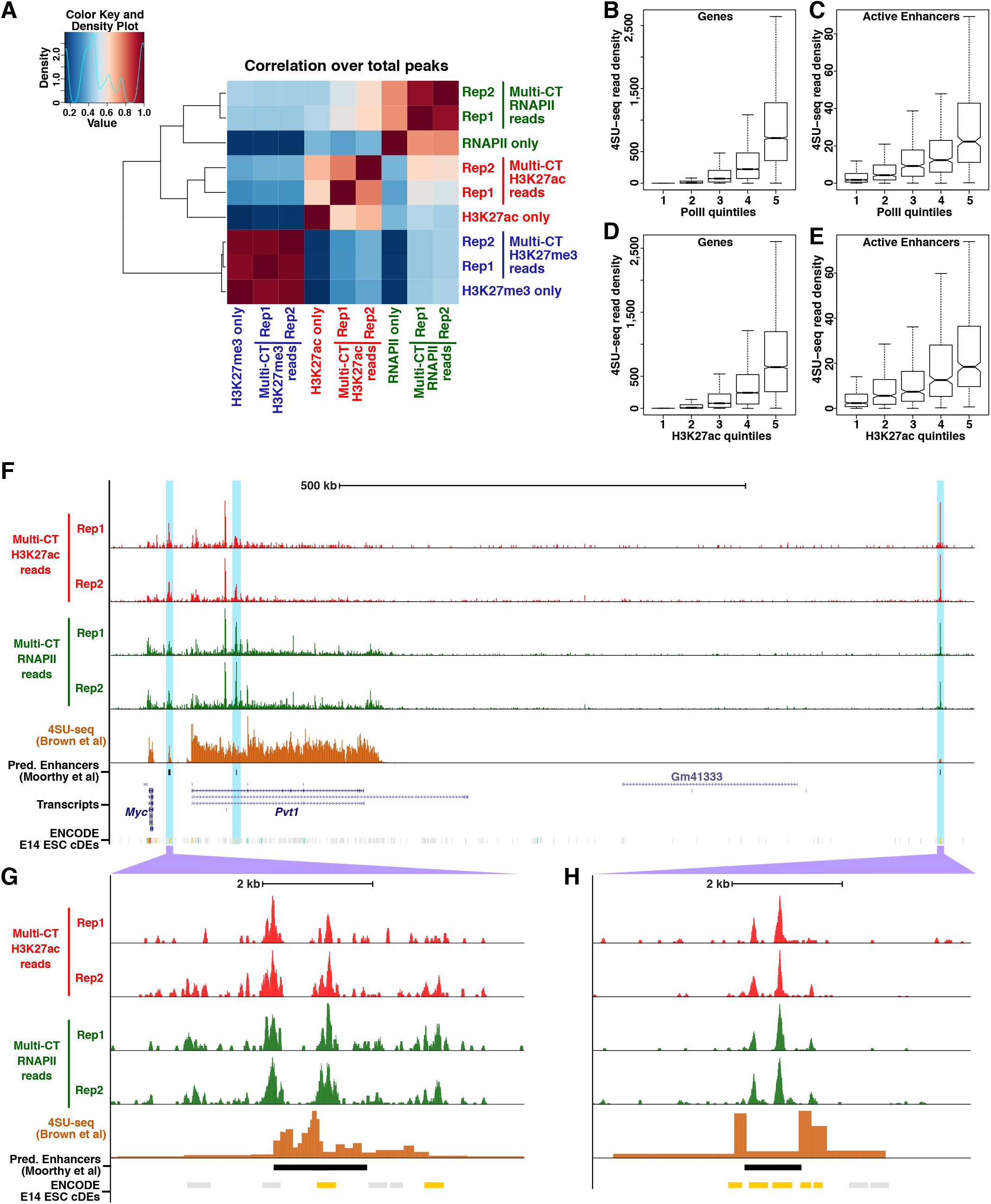
Specificity of multi-CUT&Tag profiles. **A,** Correlation matrix of single and triple Ab multi-CUT&Tag maps. Pearson correlations were calculated using the normalized read counts surrounding total H3K27ac, H3K27me3, and RNAPII peaks from multi-CUT&Tag libraries. **B-E,** Nascent transcription levels, as measured by 4SU-seq (GSE93538) for genes (B, D) or predicted ESC CREs (C, E) separated into quintiles based on read densities from multi-CUT&Tag maps of RNAPII (B, C) or H3K27ac (D, E). Predicted CREs were taken from a previous study (Moorthy et al., 2016). **F,** Browser tracks of multi-CUT&Tag maps of H3K27ac, RNAPII, and 4SU-seq depicting a region downstream of the Myc gene enriched for predicted cell type-specific enhancers. **G-H**, Detailed maps of two candidate CREs marked by H3K27ac and RNAPII, that exhibit nascent transcription (4SU-seq) found at many enhancers. ENCODE candidate CREs (cCREs) from E14 ESCs are shown for reference; potential promoter-distal enhancers in this dataset are highlighted in yellow.

Active enhancers and promoters are frequently marked by H3K27ac, RNAPII, and other activators of transcription. To examine the utility of multi-CUT&Tag maps to identify actively transcribed regions, we stratified genes or previously identified candidate ESC enhancers (Moorthy et al., 2016) by H3K27ac or RNAPII read density from multi-CUT&Tag maps and quantified nascent transcription from a published mESC 4SU-seq dataset (Brown et al., 2017) at each stratum of H3K27ac or RNAPII. As expected, higher levels of H3K27ac and RNAPII accompanied higher levels of nascent transcription, both at genes and candidate enhancers (Figure 2B-E). As one example, we observed strong peaks of enrichment for both H3K27ac and RNAPII at several predicted enhancers downstream of the *Myc* gene, including some located within actively transcribed genes marked by broad domains of moderate H3K27ac and RNAPII read density (Figure 2F-H). These data suggest multi-CUT&Tag is an effective tool for identification of candidate CREs, enabling different enhancer/promoter marks to be mapped in a single experiment, which drastically reduces the numbers of cells and experiments needed for this purpose.

### Co-localization of different chromatin proteins identified by multi-CUT&Tag

Numerous TFs, histone modifications, and chromatin regulatory proteins are thought to colocalize on chromatin, as inferred by the overlap of their ChIP-seq maps. In particular, large-scale mapping efforts have shown co-localization of dozens of proteins and epigenetic marks to known or predicted CREs (Consortium et al., 2020; Ho et al., 2014). However, the overlap among multiple chromatin proteins revealed in these and other studies could also reflect alternative binding states, where one factor or the other (but not both) bind at each overlapping site, since different factors are mapped in different samples. Consequently, a method that directly and efficiently measures co-binding of factors in the same cells is necessary to dissect the mechanisms by which CREs are utilized to direct gene expression.

Although the majority of multi-CUT&Tag reads have “homogeneous” barcodes, where barcodes corresponding to one Ab are present at both ends, a fraction of multi-CUT&Tag reads have “mixed” barcodes, with different Ab barcodes at each end (Figure 3A). Since each read is derived from a single chromosomal fragment in a single cell, reads with mixed barcodes reflect co-localization of both epitopes at the same location on the same chromosomal copy. Therefore, multi-CUT&Tag can, in principle, map the co-localization of two or more histone modifications or different chromatin proteins in cells without the requirement for single cell profiling.

**Figure 3:**
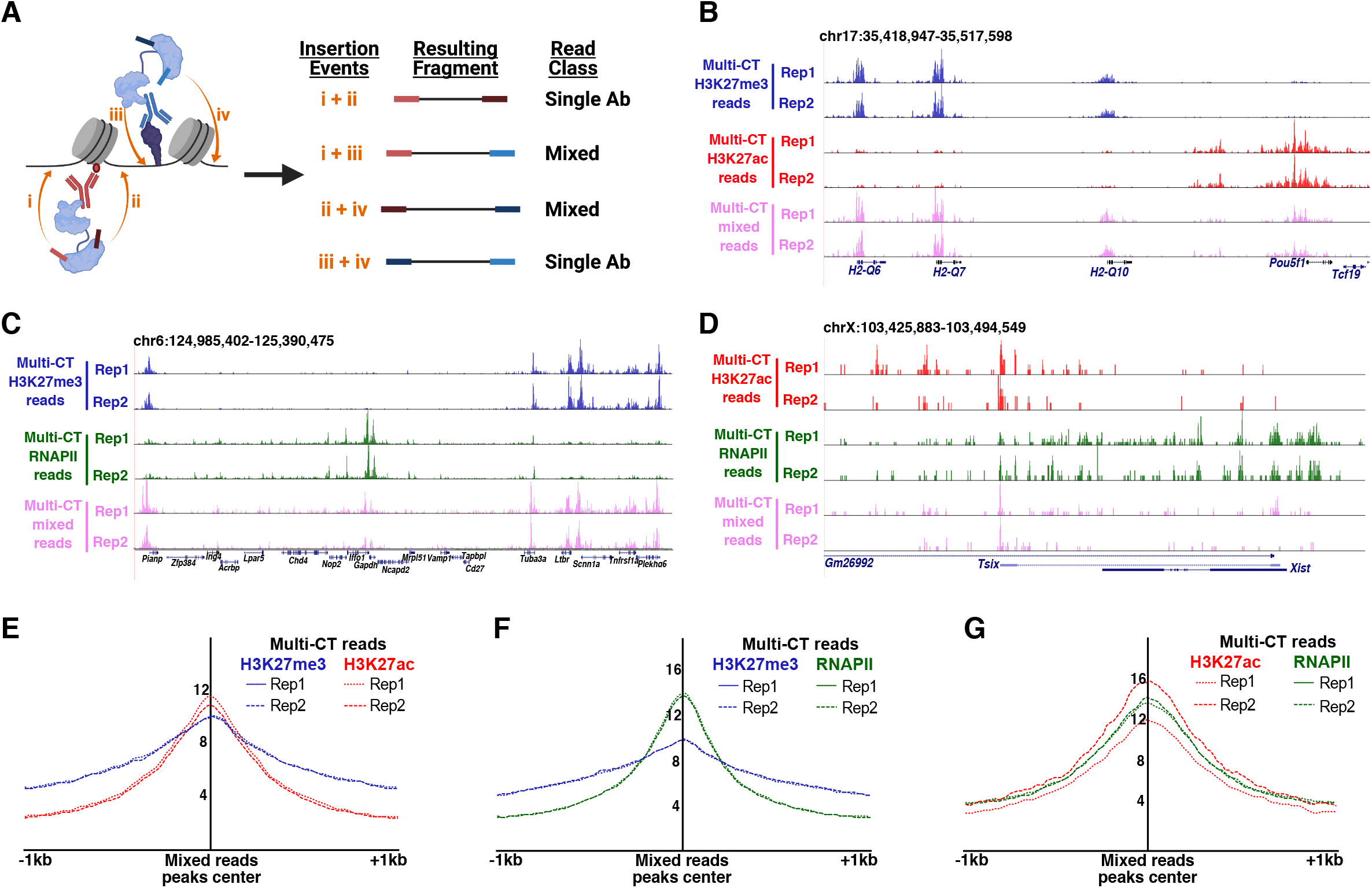
Co-localization of histone modifications and RNAPII directly detected by multi-CUT&Tag. **A,** Schematic representation of homogenous and mixed reads derived from multi-CUT&Tag **B-D,** Genome browser tracks showing enrichment of homogenous reads (with barcodes for a single Ab on both ends) and mixed reads (one barcode specific to each Ab on either end) from dual Ab multi-CUT&Tag profiles of H3K27me3+H3K27ac (B), H3K27me3+RNAPII (C), and H3K27ac+RNAPII (D). Each track is normalized to the same number of reads to allow direct comparison of homogeneous and mixed read profiles **E-G,** Aggregate enrichment of homogenous reads for each Ab listed over the peaks called from mixed reads in dual Ab multi-CUT&Tag for H3K27me3+H3K27ac (E), H3K27me3+RNAPII (F), and H3K27ac+RNAPII (G).

To evaluate the features and specificity of mixed reads from multi-CUT&Tag libraries, we compared maps of mixed reads corresponding to each pairwise combination of Abs to their homogeneous counterparts. For each pair of Abs examined, we observed enrichment of mixed reads at locations where homogeneous reads for both Abs were present (Figure 3B-D), although mixed reads were substantially less abundant than each set of homogeneous reads. Overall, 18-20% of the unique reads from all combinations of Abs were classified as mixed reads in the double Ab multi-CUT&Tag libraries. To examine whether mixed reads represented a special class of inserts within multi-CUT&Tag libraries (e.g., reads at the borders of nonoverlapping chromatin domains that may be separated by variable genomic distances), we compared the size distribution of homogenous and mixed reads. We found the size classes of each set to be nearly identical, with large populations of both nucleosome-sized and subnucleosome-sized inserts, suggesting that mixed reads did not represent a unique sub-class of inserts within the libraries (Figure S3). Finally, we observed no offset in the overall locations of mixed and homogeneous reads (Figure 3E-G), confirming that the same genomic regions produce homogeneous reads (specific for either of two overlapping epitopes) and mixed reads from cells within the same population. These data demonstrate multi-CUT&Tag can effectively map co-localization of different chromatin proteins or histone modifications within cells, even when co-localization is relatively infrequent, as is the case for H3K27me3 vis-à-vis H3K27ac or RNAPII.

### multi-CUT&Tag profiling in single cells

We and others have previously demonstrated the ability of methods adapted from CUT&RUN or CUT&Tag to map TFs and histone modifications in single cells (Bartlett et al., 2021; Bartosovic et al., 2020; Carter et al., 2019; Hainer et al., 2019; Kaya-Okur et al., 2019; Skene and Henikoff, 2017; Wang et al., 2019; Wu et al., 2020; Xiong et al., 2020; Zhu et al., 2021). Critically, in single cell protocols that utilize transposition by Tn5, many of the enzymatic steps and washes can be performed in bulk cell populations, followed by single cell isolation and barcoding using any of several different methods (Bartlett et al., 2021; Bartosovic et al., 2020; Kaya-Okur et al., 2019; Wu et al., 2020). Based in part on these previous advances, we next developed a simple strategy to adapt multi-CUT&Tag to profile multiple chromatin factors in high throughput within the same single cells.

For single-cell multi-CUT&Tag (scMulti-CUT&Tag), all steps prior to tagmentation were performed in a manner similar to bulk multi-CUT&Tag, except cells were not immobilized on concanavalin A magnetic beads. After the addition of magnesium to activate Tn5, tagmented single cells were isolated in microdroplets with gel beads containing cell-specific barcoded oligonucleotides using the 10X Genomics Chromium platform (Figure 4A). After addition of cellspecific barcodes, fragments from single cells were combined and built into scMulti-CUT&Tag libraries as described in the Methods. Each read in the resulting libraries featured cell-specific barcodes, sample barcodes, and antibody-specific barcodes surrounding the genomic DNA insert (Figure S4A).

**Figure 4:**
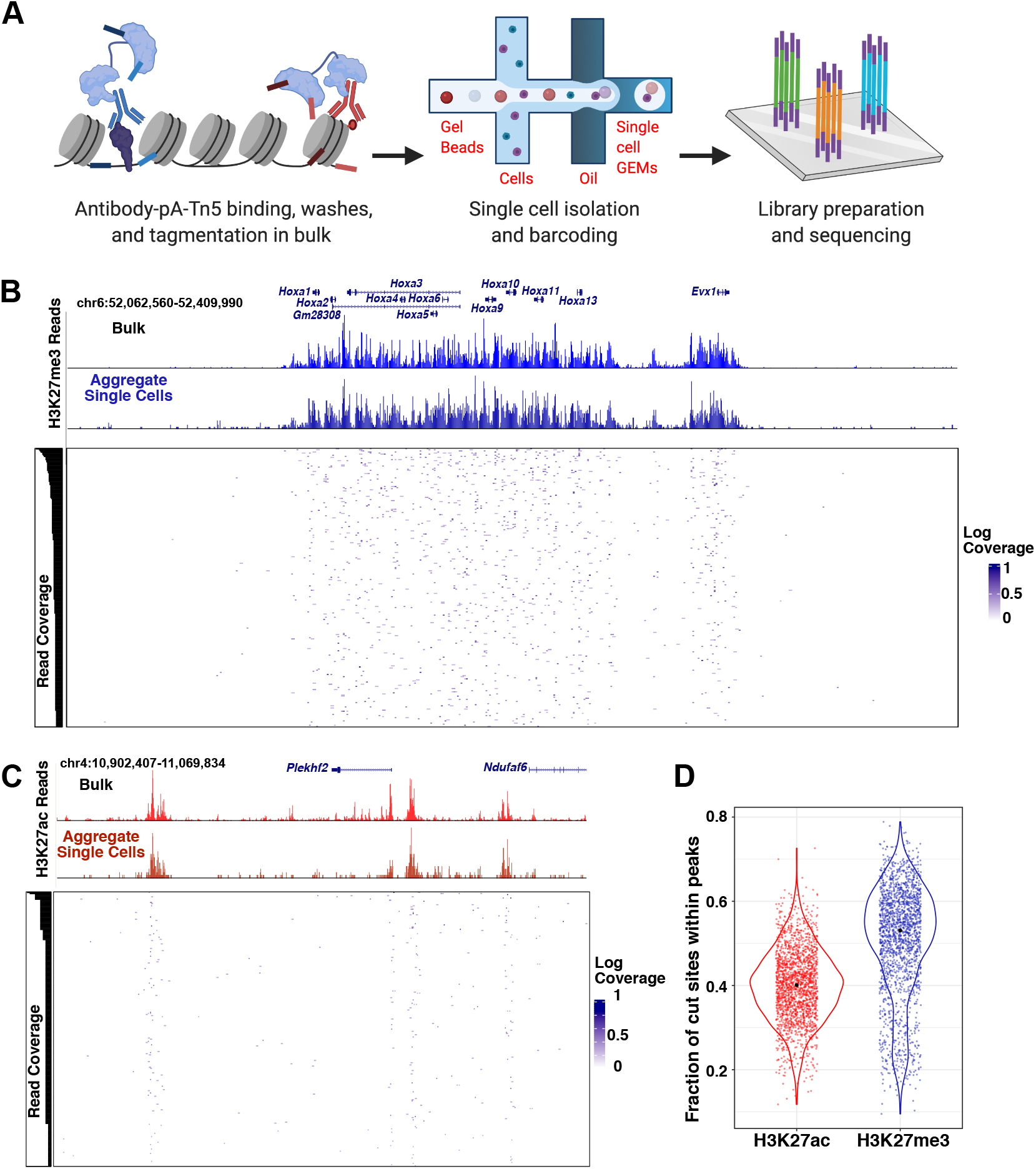
multi-CUT&Tag profiling in single cells. **A,** Diagrammatic representation of scMulti-CUT&Tag approach. **B-C,** Chromatin landscapes showing comparing bulk CUT&Tag maps with scMulti-CUT&Tag maps, both in aggregate over all single cells and individual cells, at regions of enrichment of H3K27me3 (B) and H3K27ac (C). For H3K27me3, the cells with six or more cut sites within the region were shown and for H3K27ac, the cells with at least one cut site within the region were shown. For both Abs, cells were ordered by read coverage within the regions depicted. **D,** Violin plots depicting the normalized cut sites for H3K27me3 and H3K27ac within peaks specific for specified genomic regions across single cells. Peaks were called from aggregate single cell datasets for each Ab, as described in Methods.

As proof-of-concept, we profiled H3K27me3 and H3K27ac in a mixed population of mESCs and mouse trophoblast stem cells (mTSCs). We used a custom analysis pipeline (described in Methods) to separate reads according to cell and Ab barcodes, as well as remove duplicates and quantify cut sites per cell for each antibody. First, to ensure that single-cell isolation and library preparation maintained the high specificity observed in bulk multi-CUT&Tag, we examined the aggregated reads from all cells for each antibody. Aggregate single cell maps were similar to profiles from bulk single antibody CUT&Tag for both H3K27me3 and H3K27ac at many different loci, with similarly high signal-to-noise (Figure 4B-C, Figure S4B). Next, we examined the distribution of cut sites per cell within peaks called from bulk single cells. The median percentage of cut sites within peaks was 53% for H3K27me3 and 40% for H3K27ac, with greater than 96% of cells exhibiting above 20% of their cut sites within peaks for both antibodies (Figure 4D), confirming the high specificity of scMulti-CUT&Tag.

### Pseudo-bulk analysis of scMulti-CUT&Tag profiles uncovers cell type-specific chromatin domains

Given the high fractions of informative reads in scMulti-CUT&Tag libraries (Figure 4D), we examined whether unbiased clustering could be performed to identify distinct cell types from the mixture of ESCs and TSCs profiled. To this end, we performed latent semantic indexing (LSI), followed by clustering of cells on the first two components. We first tested several thresholds of minimum reads per cell to determine whether separable clusters of cells could be identified and the number of reads per cell needed to generate informative clusters. Significantly, we found that relatively few unique reads per cell were required to produce two distinct clusters (Figure S5A). We observed excellent separation of clusters of at a threshold of 200 reads per cell (Figure 5A), with 1,949 cells meeting this threshold.

**Figure 5:**
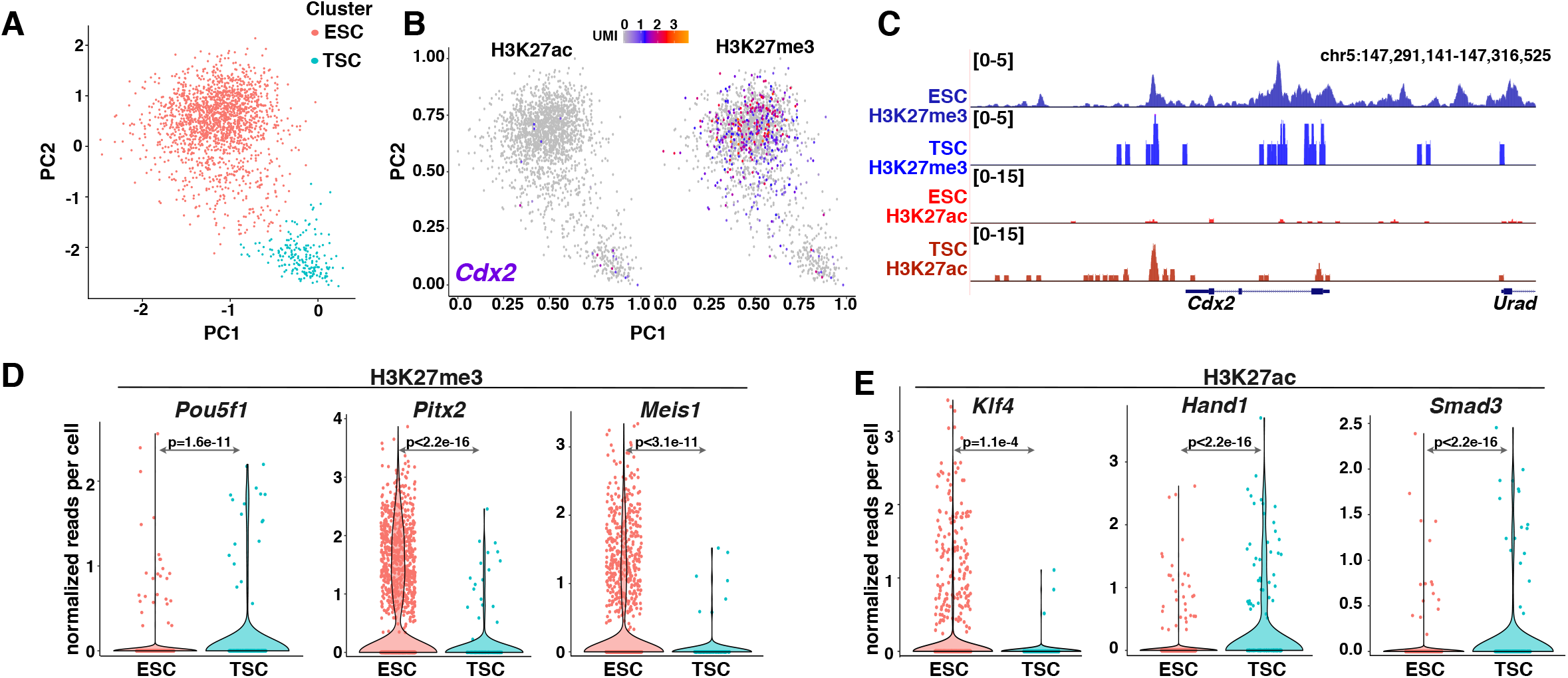
Unbiased clustering of integrated scMulti-CUT&Tag profiles uncovers cell typespecific differences in chromatin architecture. **A,** Latent semantic indexing (LSI) plots derived from integrated H3K27ac and H3K27me3 scMulti-CUT&Tag profiles, revealing cell type-specific clusters. **B-C,** LSI projections (B) and pseudo-bulk analysis (C) of H3K27me3 and H3K27ac enrichment surrounding the TSC gene *Cdx2* within each cell cluster. **D-E,** Violin plots showing mean enrichment of H3K27me3 (D) and H3K27ac (E) in ESCs and TSCs near genes exhibiting cell type-specific enrichment of one or both chromatin marks, as indicated.

Next, we generated pseudo-bulk maps of histone marks from each cluster of cells to identify which cell type made up each cluster and determine whether known features of ESC and TSC chromatin structure can be identified from scMulti-CUT&Tag maps. Clusters corresponding to ESC or TSC identities were evident based on differences in chromatin profiles at several cell type-specific marker genes. As one example, the TSC master regulatory gene *Cdx2*, which is expressed specifically in TSCs, exhibits high H3K27me3 throughout its upstream regulatory regions and gene body in ESCs and higher H3K27ac in TSCs, as expected (Figure 5B, C). To further explore differences in chromatin architecture in the ESC and TSC clusters, we identified regions of differential enrichment of both histone modifications between the two clusters. Among regions with significant cell type-specific differences in chromatin architecture, we observed regulatory regions or coding sequences of multiple genes with established roles in ESCs or TSCs (Figure 5D, E). As one prominent example, the *Hand1* gene, which encodes a transcription factor that functions in TSC differentiation, was associated with notably higher levels of H3K27ac in the TSC cluster. Additional examples include *Klf4* and *Pou5f1*, encoding key ESC transcription factors, the developmental transcription factor *Meis1*, and other lineage specific genes (Figure 5D, E). As independent validation, we observed that peaks of H3K27ac or H3K27me3 enrichment specific to the ESC cell cluster were strongly enriched in publicly available ChIP-seq maps from ESCs relative to TSCs (Chuong et al., 2013; Dunham et al., 2012) (Figure S5B). In sum, these studies confirm the ability of scMulti-CUT&Tag maps to identify different cell types from a mixed population of cells and uncover cell type-specific differences in chromatin structure.

## Discussion

Here we have described multi-CUT&Tag, an approach based on the CUT&Tag method for genome-wide chromatin mapping using targeted recruitment of pA-Tn5 to mark the genomic locations of chromatin proteins in cells (Kaya-Okur et al., 2019). This rapidly evolving technology has recently been modified multiple times for different single-cell protocols such as droplet-based cell isolation or split-pool barcoding, linear amplification of inserts, and profiling of RNA from the same cells used for mapping of chromatin proteins (Bartlett et al., 2021; Bartosovic et al., 2020; Wu et al., 2020; Xiong et al., 2020; Zhu et al., 2021). However, each of these methods can be performed on only one protein at a time per cell or pool of cells. In contrast, we use pre-coupling of pA-Tn5 to Abs to generate uniquely barcoded Ab•pA-Tn5 complexes that can be used simultaneously in the same cells to map multiple proteins. We demonstrate simultaneous use of three Ab•pA-Tn5 complexes in bulk samples and two Ab•pA-Tn5 complexes in single cells. However, the number of different Ab•pA-Tn5 combinations that can simultaneously be profiled may be significantly higher, depending on the specific combinations of epitopes to be mapped.

Multi-CUT&Tag was designed to provide two advances relative to current approaches, which will enable studies that have, until now, been unfeasible. First, epigenomic maps of multiple chromatin proteins can be generated from the same biological samples. When samples are limiting, as is the case with many human tissue specimens, sorted populations of rare cell types, or embryonic stages of various animal models, there may not be material sufficient to map multiple epitopes using standard approaches. A second advantage is the ability to directly measure co-association of different chromatin proteins. Although co-association of chromatin proteins is often inferred when two proteins have overlapping peaks of enrichment, the possibility each protein binds overlapping regions in different populations of cells cannot be ruled out when each protein is mapped separately. A significant minority of inserts from two and three Ab multi-CUT&Tag libraries contained barcodes for different Abs at each end, demonstrating co-localization of chromatin proteins can be directly measured by multi-CUT&Tag, even at the relatively low sequencing depth required for CUT&RUN and CUT&Tag based approaches (Kaya-Okur et al., 2019; Skene and Henikoff, 2017). It is worth noting that many potential co-localization studies can be performed using multi-CUT&Tag without the need for single-cell isolation and barcoding, since each mixed read reflects colocalization of two epitopes in a single cell, regardless of the number of cells used for mapping. This can be advantageous over scMulti-CUT&Tag in many cases, due to its lower cost.

We observed the expected co-localization of the active histone modification H3K27ac with RNAPII, as well as some unexpected overlap between H3K27ac and the repressive mark, H3K27me3. Although other combinations of activating/repressive marks, such as H3K4me3 and H3K27me3, are well known to co-localize at “poised” regulatory regions (Azuara et al., 2006; Bernstein et al., 2006), co-localization of H3K27ac and H3K27me3 is less well-established. Overlap between individual H3K27ac and H3K27me3 ChIP-seq maps can be observed, typically at low levels that may not be classified as peaks. Such low-level overlap from ChIP-seq maps may suggest these marks are present at the same sites in different populations of cells where chromatin structure is not homogeneously active or inactive. By mapping both marks simultaneously in the same cells, we were able to address this possibility directly, showing that these marks do indeed co-localize at some loci.

In addition to joint profiling of multiple epitopes in bulk, we also demonstrate the utility of multi-CUT&Tag in single cells. For single cell mapping, we adapted multi-CUT&Tag for use with the 10X Genomics Chromium platform, due to its ease of use and relative availability. However, we anticipate that this approach can easily be adapted to other single-cell workflows, such as the ICELL8 system or split-pool barcoding, which have previously been used in CUT&Tag studies (Bartlett et al., 2021; Kaya-Okur et al., 2019; Xiong et al., 2020; Zhu et al., 2021). Single-cell libraries exhibited similarly high signal-to-noise as bulk multi-CUT&Tag libraries and traditional CUT&Tag approaches, as determined by the fraction of cut sites within peaks. This high specificity—evident in both single-cell and pseudo-bulk browser tracks of both H3K27me3 and H3K27ac—enables clustering of cell types and identification of cell typespecific chromatin domains.

In sum, multi-CUT&Tag enables high-efficiency, simultaneous profiling of multiple chromatin proteins in the same cells. This approach will facilitate studies of co-binding of chromatin proteins that have, until now, been impractical. In addition, single-cell adaptations of multi-CUT&Tag enable profiling of different cell types within heterogeneous populations to uncover cell type-specific differences in chromatin structure.

## Limitations

The efficiency of multi-CUT&Tag is likely limited by the fact that only two Ab barcodes can be present in each unique genomic insert. Accordingly, if a very high number of different Ab•pA-Tn5 combinations are used, Ab•pA-Tn5 complexes corresponding to co-localizing chromatin proteins will likely interfere with each other by competing for insertion into the same DNA fragments. The ability of multi-CUT&Tag to directly detect co-binding of two proteins, and/or colocalization of different histone modifications, represents a major advance that enables mechanistic studies of binding cooperativity that were previously limited to biochemical approaches. However, it is difficult to distinguish whether mixed multi-CUT&Tag reads for histone modifications exist on the same nucleosome or adjacent nucleosomes, since adapters mapping between two nucleosomes could be derived from pA-Tn5 complexes recruited by histone modifications on either nucleosome. This problem may be mitigated in the case of sequence-specific DNA binding proteins, where regions with well-separated sets of binding sites for each protein can be examined.

One advantage of all CUT&Tag-based approaches is their high specificity, as determined by the fractions of reads within peaks compared with ChIP-seq approaches, enabling generation of high-quality maps from as few as 2-3 million reads (Kaya-Okur et al., 2019, 2020). The flip side to this advantage is that fewer unique inserts per cell are obtained from scMulti-CUT&Tag compared with alternative epigenetic profiling approaches such as scATAC-seq, as previously demonstrated for standard CUT&Tag and related techniques (Bartosovic et al., 2020; Carter et al., 2019; Kaya-Okur et al., 2019; Wang et al., 2019; Wu et al., 2020; Zhu et al., 2021). While a majority of reads from bulk ChIP-seq and scChIP-seq maps fall outside of peaks, as detailed by others (Kaya-Okur et al., 2020), these “background” reads are less abundant in bulk and single cell CUT&Tag/multi-CUT&Tag maps, reducing the number reads obtained per cell. Despite this property, we found that aggregate scMulti-CUT&Tag data from ~3,500 single cells faithfully recapitulated bulk multi-CUT&Tag maps. We further showed that a mixed population of ESCs and TSCs could easily be segregated on the basis of their joint H3K27ac/H3K27me3 profiles, demonstrating the power of the scMulti-CUT&Tag approach. Interestingly, TIP-seq, a recent modification of CUT&Tag, appears to significantly increase the numbers of unique reads per cell in single-cell studies by virtue of a linearly amplified RNA readout (Bartlett et al., 2021). Therefore, adaptation of the TIP-seq RNA amplification strategy into scMulti-CUT&Tag may further enhance this method.

## Acknowledgements

We thank Sundeep Kalantry for providing TSCs, Pranitha Vangala for advice on single cell analyses, and Jennifer Benanti for critical comments on the manuscript. This work was supported by NIH grants R01HD072122 and R01HD093783 to T.G.F. and R21CA236594 to M.G.

## Author contributions

S.G. and T.G.F. designed the study. S.G. and N.W.H. performed preliminary method development studies. S.G. optimized the multi-CUT&Tag approach and performed all experiments included in manuscript. Bulk data were analyzed by S.G. and T.G.F., and single cell data were analyzed by Y.W. and M.G. Finally, S.G. and T.G.F. wrote the manuscript with input from all authors.

## Experimental model and subject details

### Cell culture

E14 mouse embryonic stem cells (ESCs) were grown DMEM-high glucose media (MilliporeSigma, D6546-500ML) containing 10% Fetal Bovine Serum (MilliporeSigma, F2442), 2 mM L-Glutamine (Corning, 25-005-CI), MEM Nonessential Amino Acids (Corning, 25-025-CI), β-mercaptoethanol (MilliporeSigma, M6250) and recombinant leukemia inhibitory factor. 5-4 mouse trophoblast stem cells (TSCs) were grown in RPMI 1640 media (ThermoFisher, 11875093) supplemented with 20% Fetal Bovine Serum, 2 mM L-Glutamine, 100 μM β-mercaptoethanol containing 25 ng/mL FGF4 (Peprotech, 100-31) and 1 μg/mL heparin (Sigma, H3149). 70% of the the TSC media was pre-conditioned by growing on feeder cells before use. Both ESCs and TSCs were maintained on plates coated with 0.2% gelatin.

## Method details

### pA-Tn5 purification and antibody coupling

Histidine tagged pA-Tn5 was produced by Gibson cloning gBlocks (IDT) encoding 6-histidine tagged protein A fused to Tn5 transposase through a flexible linker (as described (Kaya-Okur et al., 2019)) into pET28a cut with NcoI and BamHI. Protein was expressed overnight at 18 °C as described (Kaya-Okur et al., 2019) in BL21 DE3 pLysS bacteria and purified at 4 °C using TALON beads according to the manufacturers’ recommendations. After dialysis into pA-Tn5 storage buffer (20mM HEPES pH 7.2, 0.2M NaCl, 0.2mM EDTA, 2mM DTT, 0.2% TritonX-100, 50% glycerol), we decided to remove trace impurities as follows. Protein was diluted four-fold into HN_50_TE (20mM HEPES pH 7.2, 10mM NaCl, 0.2% TritonX-100, 0.5mM EDTA). Diluted protein was bound to Q Sepharose Fast Flow in batch, and the unbound protein was bound to SP-Sepharose Fast Flow. After washing with approximately 20 bed volumes of HN_50_TE and 10 bed volumes of HN_150_TE (same as HN_50_TE except 150mM NaCl), pA-Tn5 was eluted twice with three bed volumes of HN_800_TE. Eluate was dialyzed into pA-Tn5 storage buffer (see above). Aliquots were quick frozen and stored long-term at −80 °C, and short term at −20 °C.

25 μL of purified pA-Tn5 (21μM) was loaded with barcoded Tn5 adapter oligos (Sup. Table 1), with sequences described previously (Amini et al., 2014), by incubating with 35 μL of 100% glycerol and 35 μL of 45 μM equimolar mixture of annealed Tn5 adapters A and B. Adapter loaded pA-Tn5 were incubated with the antibody of interest (at approximately 2:1 ratio of pA-Tn5:antibody) for 4 hours or overnight at 4 °C. To purify the antibody-pA-Tn5 complex, His-Tag purification was performed using Dynabeads His-Tag Isolation and Pulldown (Invitrogen, 10103D). Beads were allowed to bind to antibody-pA-Tn5 complex for 2 hours at 4°C, washed twice with phosphate buffered saline (PBS), and the bound complexes were eluted using PBS containing 300mM imidazole. Imidazole was removed from the eluted purified complexes by buffer exchange using Amicon Ultra-0.5mL Centrifugal filters (Millipore, UFC501096). The purified antibody-pA-Tn5 complex was stored in PBS with 50% glycerol at −20 °C.

### Bulk multi-CUT&Tag and library preparation

Differentially barcoded, purified antibody-pA-Tn5 complexes were used to perform multi-CUT&Tag experiments. Mouse ESCs or TSCs were trypsinized and counted using a TC-20 cell counter (Biorad). 100,000 cells were washed and resuspended in wash buffer (20 mM HEPES pH 7.5; 150 mM NaCl; 0.5 mM Spermidine; 1× Protease inhibitor cocktail) and used for bulk multi-CUT&Tag. 10 μL of Concanavalin A coated magnetic beads (Polysciences) were activated as described (Hainer and Fazzio, 2019) and added per sample and incubated at RT for 15 min. Bead-bound cells were resuspended in 100 μL Dig-wash Buffer (20 mM HEPES pH 7.5; 150 mM NaCl; 0.5 mM Spermidine; 0.05% Digitonin; 1× Protease inhibitor cocktail) containing 2 mM EDTA and 2 μg of each differentially barcoded purified antibody-pA-Tn5 complex was added. The mixture was incubated overnight at 4 °C for antibodies to bind. Cells were washed thrice in Dig-med Buffer (20 mM HEPES, pH 7.5; 300 mM NaCl; 0.5 mM Spermidine; 0.01% Digitonin; 1× Protease inhibitor cocktail) by placing the tube on the magnet stand until the solution clears and removal of all of the liquid by pipetting. The cells were then resuspended in 300 μL Dig-med Buffer containing 10 mM MgCl2 and incubated at 37 °C for 1 h to activate tagmentation. To stop tagmentation, 10 μL of 0.5 M EDTA, 3 μL of 10% SDS and 1 μL of 20 mg/mL Proteinase K was added to each tube, which were incubated at 55 °C for 1 hour. DNA was extracted by performing one phenol:chloroform extraction followed by ethanol precipitation. The DNA pellet was resuspended in 22 μL of 10 mM Tris pH 8.

The libraries were then amplified by mixing 21 μL of DNA with 2μL each of (10 μM) barcoded i5 and i7 primers, using a different combination for each sample. 25 μL NEBNext HiFi 2× PCR Master mix (NEB) was added, and PCR was performed using the following cycling conditions: 72 °C for 5 min (gap filling); 98 °C for 30 s; 17 cycles of 98 °C for 10 s and 63 °C for 30 s; final extension at 72 °C for 1 min and hold at 4 °C. 1.1× volume of Ampure XP beads (Beckman Coulter) was incubated with libraries for 10 min room temperature to clean up the PCR mix. Bead bound DNA was purified by washing twice with 80% ethanol and eluting in 20 μL 10 mM Tris pH 8.0. The libraries were quantified by Qubit and sequenced as described below.

### Bulk multi-CUT&Tag Sequencing and data analysis

Paired-end sequencing was performed on an Illumina NextSeq 500 using custom read and index primers (Sup. Table 1). The sequencing parameters were as follows: read 1—72 cycles, read 2—72 cycles, index 1—8 cycles, index 2—8 cycles (for bulk multi-CUT&Tag) or 16 cycles (for sc multi-CUT&Tag). PhiX DNA was added to 20-30%, due to the sequence homogeneity of the initial sequencing cycles, which read through regions of the adapter that are identical in all reads. Paired-end reads from each sample were then split based on their antibody specific barcodes on both ends of the fragment using novoBarcode (http://www.novocraft.com/documentation/novobarcode/). The first 42 bases of the reads were trimmed to remove the Antibody barcodes and the bases common to all Tn5 adapter sequences. The reads were then aligned to the mouse genome (mm10) using Bowtie2 with the parameters -N 1 and -X 2000. Duplicates were removed using Picard (http://broadinstitute.github.io/picard/). Reads with low quality scores (MAPQ < 10) were removed. The remaining mapped reads were then processed using the HOMER software suite (Heinz et al., 2010). Genome browser tracks were generated using the “makeUCSCfile” command, and peaks were called using the “findPeaks” command. Heatmaps and aggregation plots were made using the “annotatePeaks” command.

GEO accession numbers for published ChIP-seq datasets to which multi-CUT&Tag datasets were compared were: H3K27me3 (Mu et al., 2018) (GSE123174), H3K27ac (GSE31039), RNAPII (Zhang et al., 2018) (GSE112113), and 4SU-seq (Brown et al., 2017) (GSE93538). The datasets were aligned to mm10 using Bowtie2, processed in Homer and peaks were called using “findPeaks” command. Predicted mESC enhancers were obtained from Moorthy et al (Moorthy et al., 2016). ENCODE candidate cis regulatory elements (cCREs) from E14 ESCs (Consortium et al., 2020) were filtered using the following criteria: DNase I z-score≥2.5, H3K27ac z-score≥2.5, H3K4me3 z-score≤2.

### Single Cell Multi-CUT&Tag

For Single Cell Multi-CUT&Tag, 200,000 cells were washed in wash buffer and resuspended in 100 μL Dig-wash Buffer containing 2 mM EDTA and combinations of differentially barcoded purified antibody-pA-Tn5 complexes. At each wash step, cells were pelleted at 600 g for 3 minutes at 4°C and the supernatant was discarded. Cells were incubated overnight with antibody-pA-Tn5 complex at 4 °C. Cells were washed thrice with Dig-wash Buffer and counted during the last wash, due to the loss of a portion of cells during the wash steps. Cells were then resuspended in Dig-med Buffer at 5000cells/μL and incubated on ice for 5 minutes. To start tagmentation, we added an equal volume of Dig-med Buffer containing 20 mM MgCl2 and incubated samples at 37 °C for 1 hour.

After one hour, 2.5 μL of the tagmentation reaction (2500 cells/μL) was used for a targeted cell recovery of 4000 cells and mixed with 2.5 μL of Diluted Nuclei Buffer, 7 μL of ATAC buffer (both from the Chromium Single Cell ATAC Reagent Kit, 10X Genomics), 3 μL of 50% glycerol and 0.5 μL of 5M NaCl. The preparation of the cell barcoding master mix, loading of the sample onto Chromium ChIP E (10X Genomics), running the sample on the 10X Chromium device, and transfer of GEMs were all performed according to the manufacturer’s protocol for single cell ATAC-seq. After preparation of GEMs, the GEM-containing tube was incubated in a thermocycler under the following conditions: 72 °C for 5 min; 98 °C for 30 s; 1 cycle of 98 °C for 10 s, 59 °C for 30 s and 72 °C for 1 min and hold at 15 °C. (Note, the use of a single amplification cycle at this step differs from the manufacturer’s recommendations for scATAC-seq, which recommends >10 cycles of linear amplification to introduce cell-specific barcodes. We found that elimination of all but one amplification cycle at this step was necessary to prevent incorporation of uninserted Tn5 adapters into libraries.) Post GEM cleanup using Dynabeads MyOne SILANE and Ampure XP beads was performed according to manufacturer’s protocol. Libraries were then constructed using sample indexing primers (Single Index Kit, 10X Genomics), performing 14 cycles of PCR according to manufacturer’s protocol. Post-PCR, we performed a double size selection using Ampure XP beads according to manufacturer’s protocol and sequenced libraries as described above.

### Single Cell Multi-CUT&Tag data analysis

BCL file conversion and demultiplexing was done using cellranger-atac/1.1.0. To eliminate the effect of errors introduced in the library preparation and sequencing processes, the reads were considered as valid only if both antibody barcodes were within the Tn5 barcode list and the cell barcode was in the whitelist provided by 10X genomics. Valid reads extraction and barcode correction were done using a custom script. Potential read-though adaptors were removed by cutadapt/1.9 (Martin, 2011). Reads were aligned to the mm10 genome with bowtie2/2.4.1, and low quality reads were removed using samtools (Li et al., 2009) with -q 30. Read pairs were considered as duplicates of the same DNA fragment if they met two criteria: their cell barcodes were identical, and they had identical start and end locations in the genome. After deduplication using a custom script, the unique cut sites were then separated according to the antibody barcodes for independent downstream process. The cut sites were extended by 100 bp on each end using bedtools/2.28.0 (Quinlan and Hall, 2010) and peaks were then called using the MACS2 (Zhang et al., 2008) ‘callpeak’ command in macs/1.4.2 with --nomodel --broad flags. Peaks with fewer than 2 cut sites per base pair per million cut sites were removed as low-quality peaks. Peaks within 3000bp were then merged. Peaks and cut sites that overlapped with the ENCODE mm10 blacklist region were removed. Peak-cell matrices for both antibodies were generated independently and then combined for the downstream analysis. After removing cell barcodes with fewer than 200 unique cut sites per cell, dimension reduction was done on regions detected in more than 20 cells with LSI implemented in Seurat/3.1.4 (Stuart et al., 2019), and unsupervised density clustering was applied on the first two LSI components. Cell identity was annotated based on the H3K27ac and H3K27me3 enrichment levels near established TSC and ESC marker genes. For plotting purposes, reads per cell were normalized to the same number, i.e. median of reads per cells. The *FindMarkers* function in Seurat was used for identification of differentially enriched regions, with test.use = ‘LR’ and latent.vars set to the unique reads per cell. To test the significance of differences in H3K27me3 or H3K27ac enrichment between ESCs and TSCs, we used a two-sided Mann–Whitney–Wilcoxon (MWW) rank sum test, owing to the non-normal distributions of the data in each group.

### Data visualization

Browser tracks were made with the UCSC genome browser (Gonzalez et al., 2021). Latent semantic indexing (LSI) plots were made with Seurat (Stuart et al., 2019). Single-cell coverage plots were made with with *ComplexHeatmap* (Gu et al., 2016) in R. Heatmaps of chromatin mapping data were made with Java Treeview, and heatmaps of correlation data were made with *gplots* in R. Schematics of multi-CUT&Tag workflows were made with BioRender.

## References

Amini, S., Pushkarev, D., Christiansen, L., Kostem, E., Royce, T., Turk, C., Pignatelli, N., Adey, A., Kitzman, J.O., Vijayan, K., et al. (2014). Haplotype-resolved whole-genome sequencing by contiguity-preserving transposition and combinatorial indexing. Nat Genet 46, 1343–1349.

Azuara, V., Perry, P., Sauer, S., Spivakov, M., Jørgensen, H.F., John, R.M., Gouti, M., Casanova, M., Warnes, G., Merkenschlager, M., et al. (2006). Chromatin signatures of pluripotent cell lines. Nat Cell Biol 8, 532–538.

Bartlett, D.A., Dileep, V., Henikoff, S., and Gilbert, D.M. (2021). High throughput genome-wide single cell protein:DNA binding site mapping by targeted insertion of promoters (TIP-seq). BioRxiv 2021.03.17.435909.

Bartosovic, M., Kabbe, M., and Castelo-Branco, G. (2020). Single-cell profiling of histone modifications in the mouse brain. Biorxiv 2020.09.02.279703.

Bernstein, B.E., Mikkelsen, T.S., Xie, X., Kamal, M., Huebert, D.J., Cuff, J., Fry, B., Meissner, A., Wernig, M., Plath, K., et al. (2006). A bivalent chromatin structure marks key developmental genes in embryonic stem cells. Cell 125, 315–326.

Brown, D.A., Cerbo, V.D., Feldmann, A., Ahn, J., Ito, S., Blackledge, N.P., Nakayama, M., McClellan, M., Dimitrova, E., Turberfield, A.H., et al. (2017). The SET1 Complex Selects Actively Transcribed Target Genes via Multivalent Interaction with CpG Island Chromatin. Cell Reports 20, 2313–2327.

Bulger, M., and Groudine, M. (2011). Functional and Mechanistic Diversity of Distal Transcription Enhancers. Cell 144, 327–339.

Carter, B., Ku, W.L., Kang, J.Y., Hu, G., Perrie, J., Tang, Q., and Zhao, K. (2019). Mapping histone modifications in low cell number and single cells using antibody-guided chromatin tagmentation (ACT-seq). Nat Commun 10, 3747.

Chuong, E.B., Rumi, M.A.K., Soares, M.J., and Baker, J.C. (2013). Endogenous retroviruses function as species-specific enhancer elements in the placenta. Nat Genet 45, 325–329.

Consortium, E.P., Moore, J.E., Purcaro, M.J., Pratt, H.E., Epstein, C.B., Shoresh, N., Adrian, J., Kawli, T., Davis, C.A., Dobin, A., et al. (2020). Expanded encyclopaedias of DNA elements in the human and mouse genomes. Nature 583, 699–710.

Dunham, I., Kundaje, A., Aldred, S.F., Collins, P.J., Davis, C.A., Doyle, F., Epstein, C.B., Frietze, S., Harrow, J., Kaul, R., et al. (2012). An integrated encyclopedia of DNA elements in the human genome. Nature 489, 57–74.

Geisberg, J.V., and Struhl, K. (2004). Analysis of Protein Co-Occupancy by Quantitative Sequential Chromatin Immunoprecipitation. Curr Protoc Mol Biology 68, 21.8.1–21.8.7.

Gilmour, D.S., and Lis, J.T. (1984). Detecting protein-DNA interactions in vivo: distribution of RNA polymerase on specific bacterial genes. Proc National Acad Sci 81, 4275–4279.

Gonzalez, J.N., Zweig, A.S., Speir, M.L., Schmelter, D., Rosenbloom, K.R., Raney, B.J., Powell, C.C., Nassar, L.R., Maulding, N.D., Lee, C.M., et al. (2021). The UCSC Genome Browser database: 2021 update. Nucleic Acids Res 49, D1046–D1057.

Gu, Z., Eils, R., and Schlesner, M. (2016). Complex heatmaps reveal patterns and correlations in multidimensional genomic data. Bioinformatics 32, 2847–2849.

Hainer, S.J., and Fazzio, T.G. (2019). High-Resolution Chromatin Profiling Using CUT&RUN. Curr Protoc Mol Biology 126, e85.

Hainer, S.J., Bošković, A., McCannell, K.N., Rando, O.J., and Fazzio, T.G. (2019). Profiling of Pluripotency Factors in Single Cells and Early Embryos. Cell 177, 1319–1329.e11.

Heinz, S., Benner, C., Spann, N., Bertolino, E., Lin, Y.C., Laslo, P., Cheng, J.X., Murre, C., Singh, H., and Glass, C.K. (2010). Simple combinations of lineage-determining transcription factors prime cis-regulatory elements required for macrophage and B cell identities. Mol Cell 38, 576–589.

Ho, J.W.K., Jung, Y.L., Liu, T., Alver, B.H., Lee, S., Ikegami, K., Sohn, K.-A., Minoda, A., Tolstorukov, M.Y., Appert, A., et al. (2014). Comparative analysis of metazoan chromatin organization. Nature 512, 449–452.

Janssens, D.H., Wu, S.J., Sarthy, J.F., Meers, M.P., Myers, C.H., Olson, J.M., Ahmad, K., and Henikoff, S. (2018). Automated in situ chromatin profiling efficiently resolves cell types and gene regulatory programs. Epigenet Chromatin 11, 74.

Kaya-Okur, H.S., Wu, S.J., Codomo, C.A., Pledger, E.S., Bryson, T.D., Henikoff, J.G., Ahmad, K., and Henikoff, S. (2019). CUT&Tag for efficient epigenomic profiling of small samples and single cells. Nat Commun 10, 1930.

Kaya-Okur, H.S., Janssens, D.H., Henikoff, J.G., Ahmad, K., and Henikoff, S. (2020). Efficient low-cost chromatin profiling with CUT&Tag. Nat Protoc 15, 3264–3283.

Kinkley, S., Helmuth, J., Polansky, J.K., Dunkel, I., Gasparoni, G., Fröhler, S., Chen, W., Walter, J., Hamann, A., and Chung, H.-R. (2016). reChIP-seq reveals widespread bivalency of H3K4me3 and H3K27me3 in CD4+ memory T cells. Nat Commun 7, 12514.

Ku, W.L., Nakamura, K., Gao, W., Cui, K., Hu, G., Tang, Q., Ni, B., and Zhao, K. (2019). Singlecell chromatin immunocleavage sequencing (scChIC-seq) to profile histone modification. Nat Methods 16, 323–325.

Landt, S.G., Marinov, G.K., Kundaje, A., Kheradpour, P., Pauli, F., Batzoglou, S., Bernstein, B.E., Bickel, P., Brown, J.B., Cayting, P., et al. (2012). ChIP-seq guidelines and practices of the ENCODE and modENCODE consortia. Genome Res 22, 1813–1831.

Li, H., Handsaker, B., Wysoker, A., Fennell, T., Ruan, J., Homer, N., Marth, G., Abecasis, G., Durbin, R., and Subgroup, 1000 Genome Project Data Processing (2009). The Sequence Alignment/Map format and SAMtools. Bioinform Oxf Engl 25, 2078–2079.

Martin, M. (2011). Cutadapt removes adapter sequences from high-throughput sequencing reads. Embnet J 17, 10–12.

Moorthy, S.D., Davidson, S., Shchuka, V.M., Singh, G., Malek-Gilani, N., Langroudi, L., Martchenko, A., So, V., Macpherson, N.N., and Mitchell, J.A. (2016). Enhancers and superenhancers have an equivalent regulatory role in embryonic stem cells through regulation of single or multiple genes. Genome Res 27, 246–258.

Mu, W., Starmer, J., Yee, D., and Magnuson, T. (2018). EZH2 variants differentially regulate polycomb repressive complex 2 in histone methylation and cell differentiation. Epigenet Chromatin 11, 71.

Ong, C.-T., and Corces, V.G. (2011). Enhancer function: new insights into the regulation of tissue-specific gene expression. Nat Rev Genet 12, 283–293.

Park, P.J. (2009). ChIP–seq: advantages and challenges of a maturing technology. Nat Rev Genet 10, 669–680.

Quinlan, A.R., and Hall, I.M. (2010). BEDTools: a flexible suite of utilities for comparing genomic features. Bioinform Oxf Engl 26, 841–842.

Skene, P.J., and Henikoff, S. (2017). An efficient targeted nuclease strategy for high-resolution mapping of DNA binding sites. Elife 6, e21856.

Stuart, T., Butler, A., Hoffman, P., Hafemeister, C., Papalexi, E., Mauck, W.M., Hao, Y., Stoeckius, M., Smibert, P., and Satija, R. (2019). Comprehensive Integration of Single-Cell Data. Cell 177, 1888–1902.e21.

Wang, Q., Xiong, H., Ai, S., Yu, X., Liu, Y., Zhang, J., and He, A. (2019). CoBATCH for High-Throughput Single-Cell Epigenomic Profiling. Mol Cell 76, 206–216.e7.

Weiner, A., Lara-Astiaso, D., Krupalnik, V., Gafni, O., David, E., Winter, D.R., Hanna, J.H., and Amit, I. (2016). Co-ChIP enables genome-wide mapping of histone mark co-occurrence at single-molecule resolution. Nat Biotechnol 34, 953–961.

Wu, S.J., Furlan, S.N., Mihalas, A.B., Kaya-Okur, H.S., Feroze, A.H., Emerson, S.N., Zheng, Y., Carson, K., Cimino, P.J., Keene, C.D., et al. (2020). Single-cell analysis of chromatin silencing programs in development and tumor progression. Biorxiv 2020.09.04.282418.

Xiong, H., Luo, Y., Wang, Q., Yu, X., and He, A. (2020). Single-cell joint detection of chromatin occupancy and transcriptome enables higher-dimensional epigenomic reconstructions. Biorxiv 2020.10.15.339226.

Zhang, T., Wei, G., Millard, C.J., Fischer, R., Konietzny, R., Kessler, B.M., Schwabe, J.W.R., and Brockdorff, N. (2018). A variant NuRD complex containing PWWP2A/B excludes MBD2/3 to regulate transcription at active genes. Nat Commun 9, 3798.

Zhang, Y., Liu, T., Meyer, C.A., Eeckhoute, J., Johnson, D.S., Bernstein, B.E., Nusbaum, C., Myers, R.M., Brown, M., Li, W., et al. (2008). Model-based analysis of ChIP-Seq (MACS). Genome Biol 9, R137.

Zhu, C., Zhang, Y., Li, Y.E., Lucero, J., Behrens, M.M., and Ren, B. (2021). Joint profiling of histone modifications and transcriptome in single cells from mouse brain. Nat Methods 18, 283–292.

