## Supplemental Information for "Simultaneous profiling of multiple chromatin proteins in the same cells"

**SUPPLEMENTARY INFORMATION**

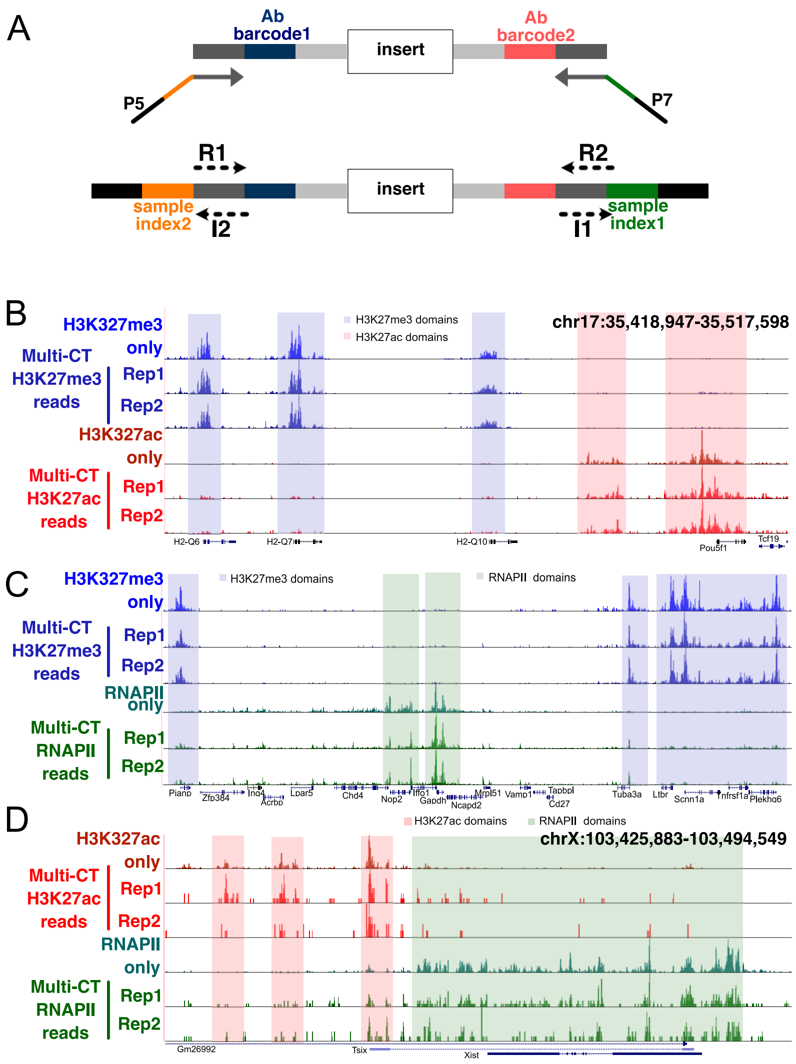


**Supplemental Figure 1: Two Ab multi-CUT&Tag profiles.**

**A,** Library construction and sequencing strategies for multi-CUT&Tag, depicting locations of Ab barcodes, sample indices, and sequencing primers for reads 1 and 2 (R1, R2) and index 1 and 2 (I1, I2) sequencing reads are shown. **B-D,** Genomic landscape showing single and double Ab multi-CUT&Tag profiles of H3K27me3+H3K27ac (B), H3K27me3+RNAPII (C), and H3K27ac+RNAPII (D). The shaded boxes represent domains enriched for the indicated epitopes.

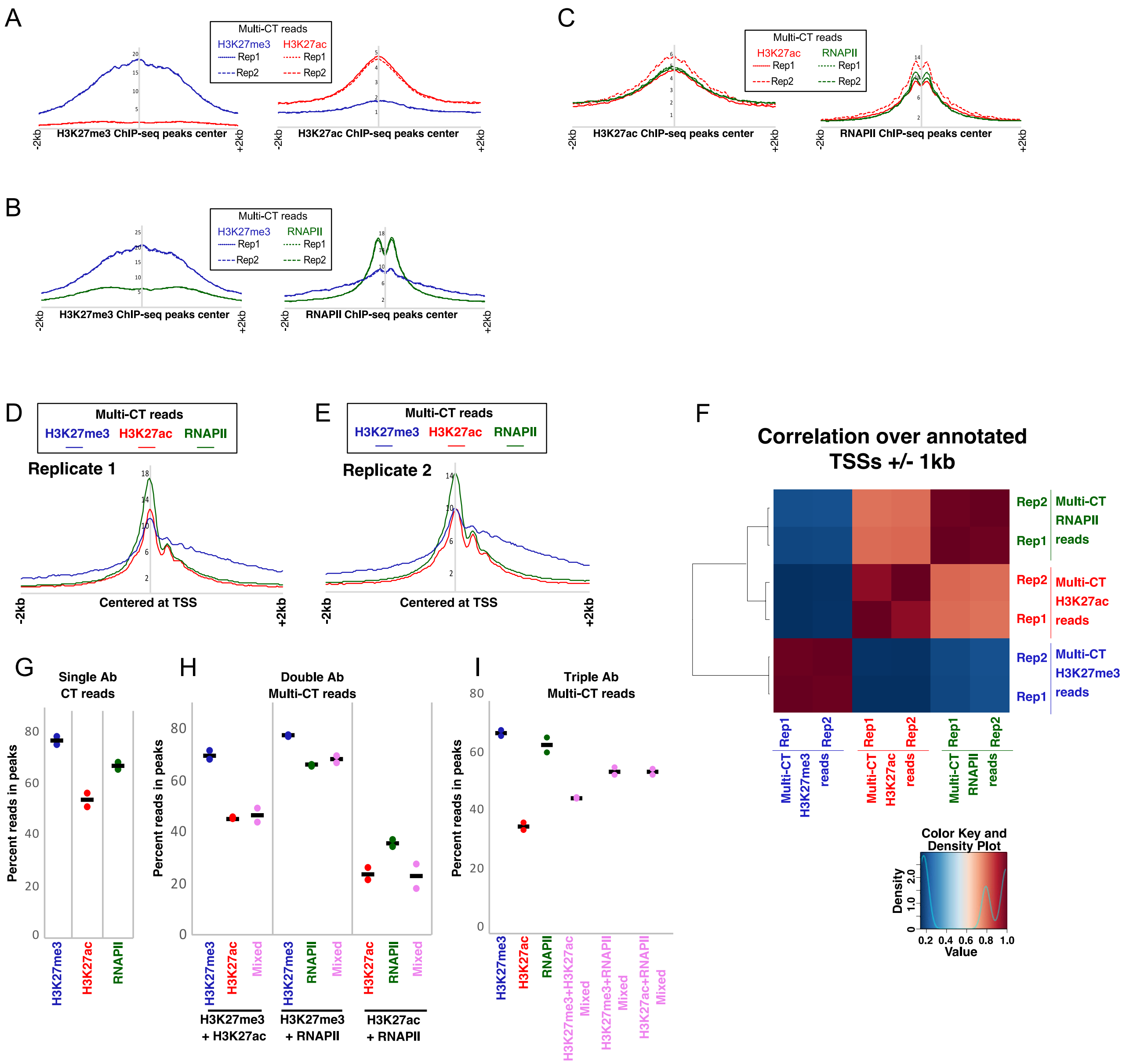


**Supplemental Figure 2: Specificity of multi-CUT&Tag maps.**

**A-C,** Average enrichment of reads from double Ab multi-CUT&Tag for H3K27me3+H3K27ac (A), H3K27me3+RNAPII (B), and H3K27ac+RNAPII (C) over published ChIP-seq peaks (as in Figure 1) corresponding to each epitope. **D-E,** Average enrichment of reads for each Ab from triple Ab multi-CUT&Tag for replicate 1 (D) and replicate 2 (E) over transcription start sites (TSS). **F,** Correlation matrix of triple Ab multi-CUT&Tag maps. Pearson correlations were calculated using the normalized read counts around annotated TSSs (+/- 1kb). **G-I,** Percent reads for H3K27me3, H3K27ac and RNAPII within peaks from single (G), double (H), and triple (I) Ab multi-CUT&Tag.

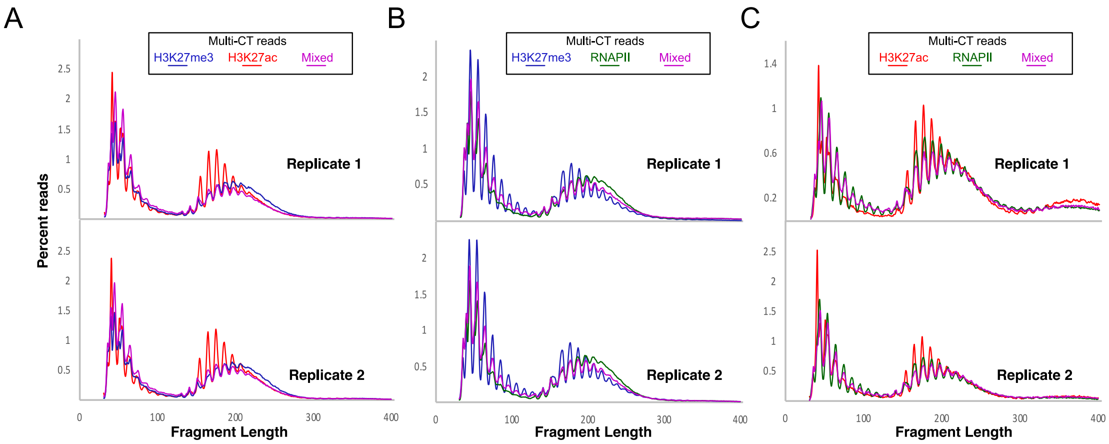


**Supplemental Figure 3: Size distribution of reads from multi-CUT&Tag.**

**A-C,** Analysis of size distribution of homogeneous and mixed reads for both replicates from double Ab multi-CUT&Tag for H3K27me3+H3K27ac (A), H3K27me3+RNAPII (B), and H3K27ac+RNAPII (C).

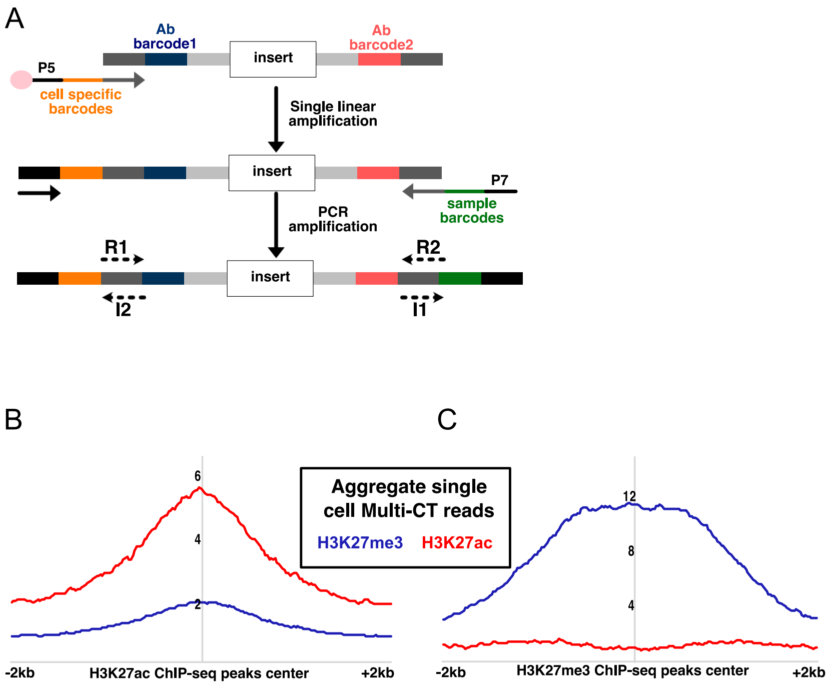


**Supplemental Figure 4: Workflow and specificity of scMulti-CUT&Tag.**

**A,** Library construction strategy for scMulti-CUT&Tag. The locations of cell specific barcodes (provided by 10X Genomics gel beads), Ab barcodes, sample barcodes, and sequencing primers for reads 1 and 2 (R1, R2) and index 1 and 2 (I1, I2) sequencing reads are shown. **B-C,** Average enrichment of reads from aggregate of scMulti-CUT&Tag over binding sites of H3K27me3 (B) and H3K227ac (C).

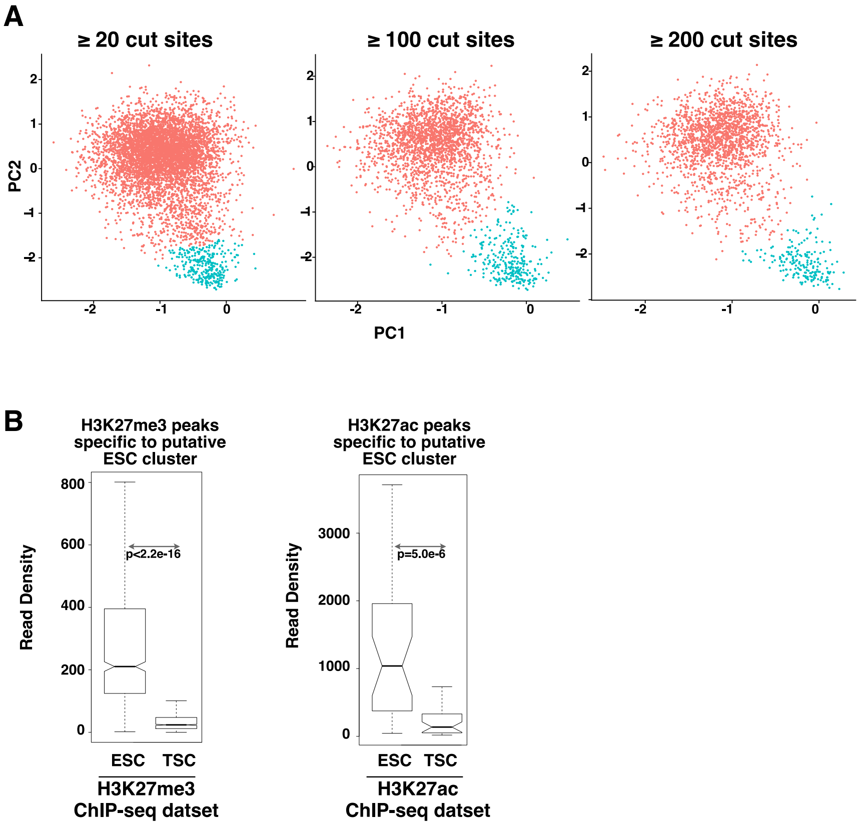


**Supplemental Figure 5: Unbiased clustering of scMulti-CUT&Tag.**

**A,** LSI plots showing cell type-specific clusters from cells with ≥20, ≥100, and ≥200 unique reads. Clusters from higher thresholds were generally similar overall to clusters using lower thresholds, but higher thresholds generally increased separation of clusters at the expense of including fewer cells. **B,** Read density from independent ESC or TSC ChIP-seq datasets within ESC-specific peaks specific to each antibody from single-cell multi-CUT&Tag.

### Supplemental Table 1. Oligonucleotide sequences used in study.

| | | |
|---|---|---|
| P5_i5_1_Universal_Connector_A | Barcoded oligo for loading to pA-Tn5 | TCGTCGGCAGCGTCTCCACGCTATAGCCTGCGATCGAGGACGGCAGATGTGTATAAGAGACAG |
| P5_i5_2_Universal_Connector_A | Barcoded oligo for loading to pA-Tn5 | TCGTCGGCAGCGTCTCCACGCATAGAGGCGCGATCGAGGACGGCAGATGTGTATAAGAGACAG |
| P5_i5_3_Universal_Connector_A | Barcoded oligo for loading to pA-Tn5 | TCGTCGGCAGCGTCTCCACGCCCTATCCTGCGATCGAGGACGGCAGATGTGTATAAGAGACAG |
| P5_i5_4_Universal_Connector_A | Barcoded oligo for loading to pA-Tn5 | TCGTCGGCAGCGTCTCCACGCGGCTCTGAGCGATCGAGGACGGCAGATGTGTATAAGAGACAG |
| P7_i7_1_Universal_Connector_B | Barcoded oligo for loading to pA-Tn5 | GTCTCGTGGGCTCGGCTGTCCCTGTCCCGAGTAATCACCGTCTCCGCCTCAGATGTGTATAAGAGACAG |
| P7_i7_2_Universal_Connector_B | Barcoded oligo for loading to pA-Tn5 | GTCTCGTGGGCTCGGCTGTCCCTGTCCTCTCCGGACACCGTCTCCGCCTCAGATGTGTATAAGAGACAG |
| P7_i7_3_Universal_Connector_B | Barcoded oligo for loading to pA-Tn5 | GTCTCGTGGGCTCGGCTGTCCCTGTCCAATGAGCGCACCGTCTCCGCCTCAGATGTGTATAAGAGACAG |
| P7_i7_4_Universal_Connector_B | Barcoded oligo for loading to pA-Tn5 | GTCTCGTGGGCTCGGCTGTCCCTGTCCGGAATCTCCACCGTCTCCGCCTCAGATGTGTATAAGAGACAG |
| Tn5MErev | Reverse Primer for loading to pA-Tn5 | [phos]CTGTCTCTTATACACATCT |
| Read1 | Custom read1 primer for multi-CUT&Tag | TCGTCGGCAGCGTCTCCACGC |
| Read2 | Custom read2 primer for multi-CUT&Tag | GTCTCGTGGGCTCGGCTGTCCCTGTCC |
| Index1 | Custom Index1 primer for multi-CUT&Tag | GGACAGGGACAGCCGAGCCCACGAGAC |
| Index2 | Custom Index2 primer for multi-CUT&Tag | GCGTGGAGACGCTGCCGACGA |
| PE read1 | Read1 for sequencing phiX | ACACTCTTTCCCTACACGACGCTCTTCCGATCT |
| PE read2 | Read2 for sequencing phiX | CGGTCTCGGCATTCCTGCTGAACCGCTCTTCCGATCT |
